# Fluorescent Glycan Fingerprinting of SARS2 Spike Proteins

**DOI:** 10.1101/2021.01.16.426965

**Authors:** Zhengliang L Wu, James M Ertelt

## Abstract

Glycosylation is the most common post-translational modification and has myriad biological functions. However, glycan analysis and research has always been a challenge. Here, we would like to present new techniques of glycan fingerprinting based on enzymatic fluorescent labeling and gel electrophoresis. The method is illustrated on SARS-2 spike (S) glycoproteins. SARS-2, a novel coronavirus and the causative agent of COVID-19 pandemic, has devastated the world since the end of 2019. To obtain the N-glycan fingerprint of a S protein, glycans released from the protein are first labeled through enzymatic incorporation of fluorophore-conjugated sialic acid or fucose, and then separated on acrylamide gel through electrophoresis, and finally visualized with a fluorescent imager. To identify the labeled glycans of a fingerprint, glycan standards and glycan ladders that are enzymatically generated are run alongside the samples as references. By comparing the mobility of a labeled glycan to that of a glycan standard, the identity of glycans maybe determined. Due to lack of enzyme for broad O-glycans releasing, O-glycans on the RBD protein are labeled with fluorescent sialic acid and digested with trypsin to obtain labeled glycan peptides that are then separated on gel. Glycan fingerprinting could serve as a quick way for global assessment of the glycosylation of a glycoprotein.

## Introduction

Glycosylation is the most common post-translational modification found on membrane and secreted proteins, thereby increase the stability and solubility of these proteins and involves in cell-cell interaction ^1, 2^. Glycans also play various roles in immunity, such as modulating the T cell activation, leukocyte homing, and the immunogenicity of proteins ^3, 4^. Autoimmune diseases are believed to be caused by aberrant glycosylation ^5^. By taking advantage of these biological functions of glycans, many viral pathogens hijack the glycosylation machinery of the host cells to camouflage their own proteins to increase the physiological properties of these proteins and evade host immune surveillance ^6-8^.

Glycan analysis is critical for researchers to better understand the biological functions of glycosylation on individual glycoproteins. Current methods for glycan analysis are mainly via mass spectrometry. However, mass spectrometry is expensive and requires highly trained expertise to perform, making it beyond the scope of many labs around the world. Besides, mass spectrometry data can have great variations, even if the data are from a same glycoprotein but generated from different labs. For example, the NIST Monoclonal Antibody Reference Material 8671 has been analyzed by 66 labs around the world and the diversity of the results was enormous, with the number of glycan compositions identified by each laboratory ranging from 4 to 48 ^9^. Therefore, alternative methods for glycan analysis that can be conveniently performed in common laboratories and can generate reproducible data are necessary.

Since the end of year 2019, the new severe acute respiratory syndrome coronavirus-2 (SARS-CoV-2) has become a major threat to human health and caused an unprecedented pandemic in history ^10^. The virus utilizes its membrane spike (S) glycoprotein to bind to human angiotensin-converting enzyme 2 (ACE-2) to initiate the invasion of host cells ^11, 12^. The S protein is proteolytically cleaved into two functional subunits, S1 and S2. S1 contains a receptor binding domain (RBD) and is responsible for the initial attachment of the virus to the surface of host cells and S2 is responsible for membrane fusion that triggers entry of the virus into the host cells ^13^. The S protein contains 22 N-glycan sequons (N-X-S/T motifs, where X is any amino acid except proline) and it is believed that glycosylation on S protein serves to camouflage the immunogenic epitopes of the protein thereby enhancing the virus’s ability to evade the host immune response ^14^. In turn, the S protein is the principal target for vaccine development and a critical component of serological assays ^15-17^. To enhance the success of vaccination and serological testing, understanding the glycosylation of the protein is critical. Previously, mass spectrometry analysis revealed that the spike protein is highly modified with complex, hybrid and oligomannose N-glycans ^14, 18, 19^; however, the glycan profiles reported by these labs varied greatly. For example, while Watanabe, Y. *et al* reported that only complex type of N-glycans were found on the RBD domain, Shajahan, A. *et al* and Zhou, D. *et al* showed that oligomannoses were the major glycans found on the RBD domain. In another example, while Shajahan, A. *et al* reported the presence of O-glycans on the RBD domain, it was not reported by Watanabe, T. *et al* and Zhou, D. *et al*. This discrepancy again necessitates alternative methods for glycan research and analysis.

Here, we report novel glycan fingerprinting techniques, in which N-glycans on a glycoprotein are first released by peptide N-glycosidase F (PNGase F) treatment and then labeled by a sialyltransferase or fucosyltransferase with a fluorophore-conjugated sialic acid or fucose ^20, 21^, and O-glycans on a glycoprotein are labeled first followed by trypsin digestion, and finally the freed labeled glycans or glycopeptides are separated by sodium dodecyl sulfate polyacrylamide gel electrophoresis (SDS-PAGE). These methods allow quick assessment of the glycosylation pattern of a glycoprotein. As an exemplification, we applied these techniques to study the glycosylation of several S protein constructs of SARS2 expressed from different host cells.

## Results

### Non-reducing End Nomenclature of N-Glycans

Previously, we showed that fluorophore-conjugated glycans can be separated on SDS-PAGE ^21^. To apply the method to glycan analysis, it is necessary to characterize those separated glycans. For convenience, we would first introduce a nomenclature for these N-glycans (Fig. 1).

**Fig. 1.**
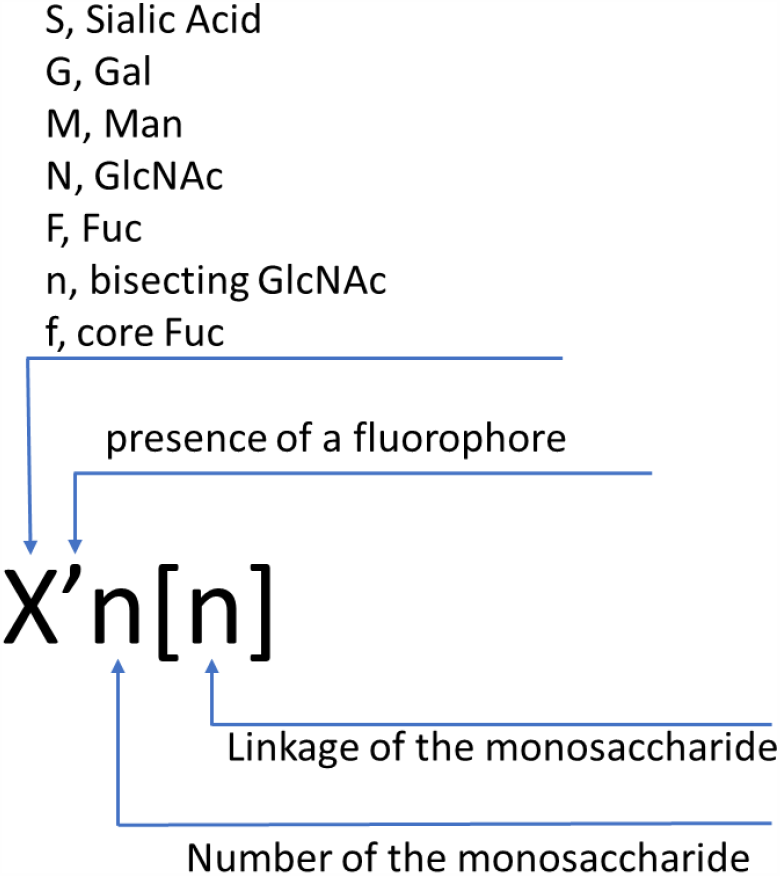
Short Name for a Non-reducing End Monosaccharide. A non-reducing end monosaccharide of an N-glycan is represented with a single capital letter followed by the number of the monosaccharide present in the glycan. If necessary, a number in a parenthesis is used to specify the linkage of the monosaccharide. A prime symbol is used when the monosaccharide is conjugated to a fluorophore. A short name of an N-glycan is a composite of all the non-reducing end monosaccharides.

In this method, only monosaccharides at the non-reducing ends of an N-glycan are specified. The rules of this method are the following. 1^st^, sialic acid, galactose, mannose, glucosamine and fucose that are commonly found at the non-reducing ends of N-glycans are represented with capital letter S, G, M, N, and F, respectively, with the exception that bisecting GlcNAc and core fucose are represented with small letter n and f, respectively. 2^nd^, a prime symbol is used to indicate if a monosaccharide is conjugated to a fluorophore. 3^rd^, the number of a non-reducing end monosaccharide of an N-glycan is specified after the letters, which is usually no more than five as this is the maximal number of the branches found on an N-glycan. 4^th^, the linkage of a non-reducing end monosaccharide is represented as a number within []. For example, S1[3]N1nf’ represents a biantennary N-glycan that contains an α,3-linked sialic acid, a GlcNAc residue with no specification on linkage, a bisecting GlcNAc and a fluorophore-conjugated core fucose at its non-reducing ends. Since there are only one bisecting GlcNAc residue and only one core fucose residue on common N-glycans, the number and linkage for these monosaccharides are omitted.

### Electrophoretic Mobility of Cy5-labeled Glycans on SDS PAGE

To correlate the glycan structures to their mobility, we established a series of labeled glycans based on a biantennary antibody glycan N2 (known as G0 in common antibody glycan nomenclature) and separated them on SDS-PAGE (Fig. 2). N2 was first labeled by FUT8 with Cy5-conjugated fucose to become N2f’. A series of glycans were then generated enzymatically based on N2f’ (Fig. 2A and 2B). N2 was also extended by B4GalT1 with or without prior modification by FUT8 and finally labeled by ST6Gal1 with Cy5-conjugated sialic acid to generate S’1[6]G1f and S’1[6]G1 (Fig. 2C). The following observations were noticed regarding the mobility change caused by the addition of different monosaccharides. First, addition of a neutral monosaccharide such as a Gal, GlcNAc, and Fuc to a glycan slows down the mobility of the glycan at a noticeable rate. Second, addition of a bisecting GlcNAc slows down the mobility of a glycan at roughly the half rate of that of a β,6-linked GlcNAc. Third, addition of a sialic acid residue significantly increases the mobility of a glycan, with even more increase on mobility by an α,6-linked sialic acid than by an α,3-linked sialic acid. Likewise, when a monosaccharide can be added at multiple positions on a glycan, intermediate glycosylation products exhibit intermediate mobilities, for example, G1N1f’ moves faster than G2f’, and S1[6]G1f’ moves slower than S2[6]f’ (Supplemental Fig. 1). Intermediate products were only observed within a short time window and were converted to final products after prolonged incubation.

**Fig. 2.**
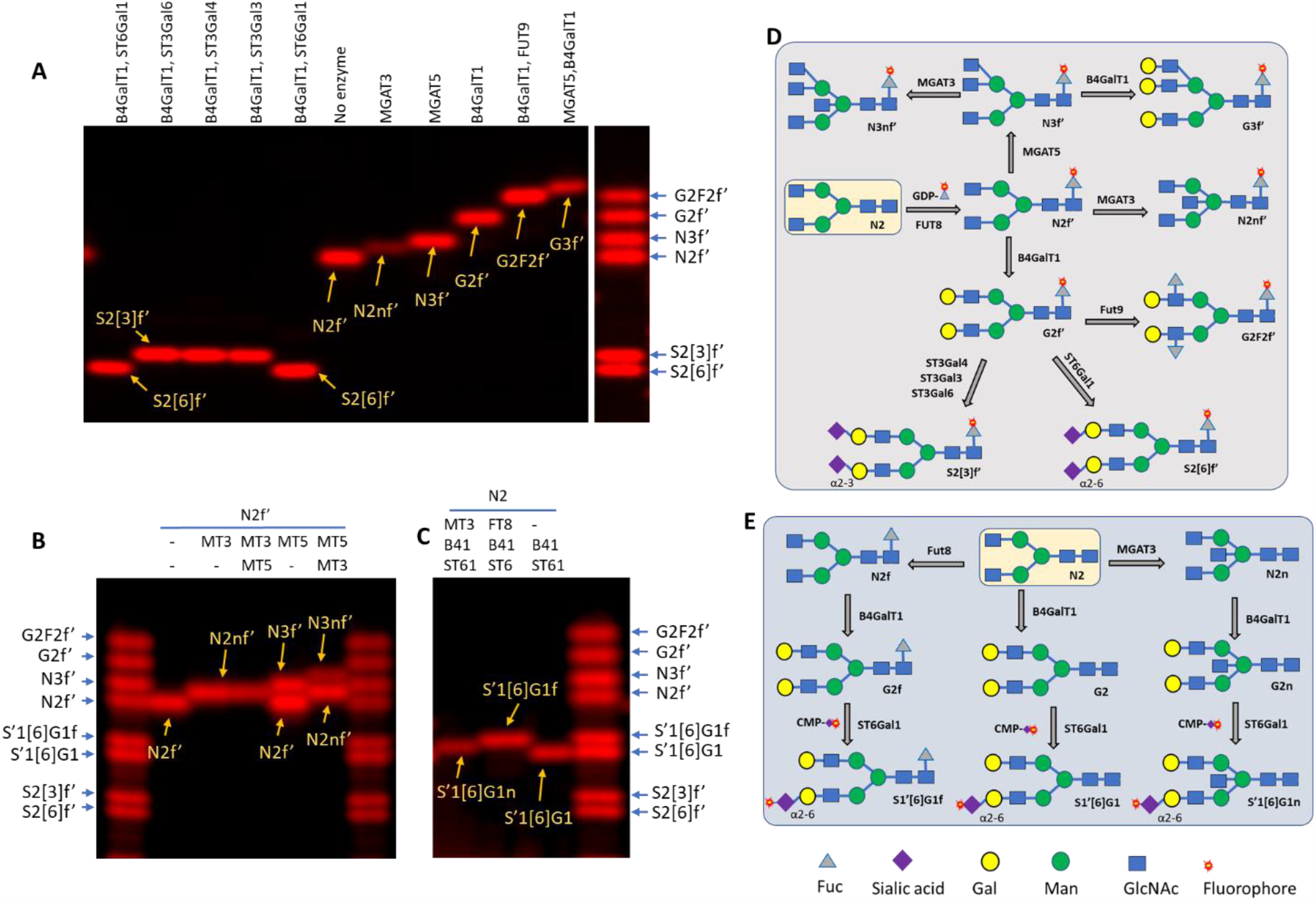
Relative mobilities of various Cy5-labeled glycans and their enzymatic synthesis. All glycans were enzymatically synthesized starting from the glycan N2 (known as G0 previously). The Glycan ladders were composed by combining some of the labeled glycans. Except enzymatic reactions of FUT8, MGAT3 and MGAT5, all other enzymatic reactions resulted in one or two intermediates as that there are two or more branches in each glycan that allow enzymatic modification (see Supplemental Fig. 1 for intermediates). The labeled glycans were separated on 17% gel and visualized with a fluorescent imager. (A) Relative mobilities of glycans that were labeled at the core-fucose. The enzymes used for generating these glycans are listed on the top of the image. (B) Relative mobility change on N2f’ by addition of a bisecting GlcNAc introduced by MGAT3 (MT3) versus a β1-6GlcNAc introduced by MGAT5 (MT5). MGAT3 and MGAT5 were introduced in different orders. The 2nd enzyme was introduced 30 minutes after the 1st enzyme. N3nf’ containing both a bisecting and a β1-6GlcNAc can only be observed when MGAT5 was introduced first. (C) Relative mobility change on S’[6]G1 by addition of a bisecting GlcNAc versus a core-fucose. The glycans were generated and labeled starting from N2 with the enzymes indicated at the top of the gel in the specified order. MT3, MGAT3; FT8, FUT8; B41, B4GalT1; ST61, ST6Gal1. (D) Schemes for enzymatic generation of the labeled glycans in (A) and (B). (E) Schemes of enzymatic generation of the labeled glycans in (C). Two fluorophore sialic acids can be introduced to each of the glycan, but only glycans with one Cy5-conjugated sialic acid were displayed in (B), (C) and (E).

### Selection of Labeling Enzyme and Optimization of Substrate Concentrations

Before we proceed to fingerprinting glycans released from various SARS2 spike proteins, we screened the labeling enzymes and optimized the substrate concentration for the labeling reaction using glycans released from the RBD protein expressed in CHO cells as the substrates. The glycans were first probed by various sialyltransferases, including ST6Gal1 that generates α2,6-sialylated N-glycans^22^, and, ST3Gal3, ST3Gal4 and ST3Gal6 that generate α2,3-sialylated N-glycans^23, 24^. Among these enzymes, ST6Gal1 and ST3Gal6 gave stronger signals (Supplemental Fig. 2A) and were chosen for the following glycan fingerprinting study. Stronger signal intensities were also observed when the substrate input was around 2 µg (Supplemental Fig. 2B) and the donor CMP-Cy5-Sialic Acid input was around 0.4 nmol (Supplemental Fig. 2C), therefore these conditions were chosen for the following fingerprinting study.

### N-Glycan Finger-printing Study of SARS-2 Spike proteins with ST6Gal1

N-glycans released from the following SARS-2 spike protein constructs with or without prior desialylation were then labeled with ST6Gal1/CMP-Cy5-Sialic Acid: RBD domain expressed in Sf21 cells (RS), RBD domain expressed in CHO cells (RC), RBD domain expressed in HEK293 cells (RH), full length spike protein expressed in CHO cells (SC), full length spike protein expressed in HEK293 cells (SH), and S1 protein expressed in HEK293 cells (S1H). As the presence of oliogomannose glycans on S proteins were reported previously ^14, 18^, FUT8/GDP-AlexaFluor555-Fuc together with MGAT1/UDP-GlcNAc that allows the labeling of Man3 and Man5 ^21^ were also added into the final labeling reactions to reveal these glycans. ST6Gal1 labeling revealed a series of bands with large variations from all constructs except RS (Fig. 3A). In general, desialylation resulted in elimination of some fast-moving bands and increased labeling on some slow-moving bands, suggesting the existence of both sialylated and asialylated glycans on these proteins.

**Figure 3.**
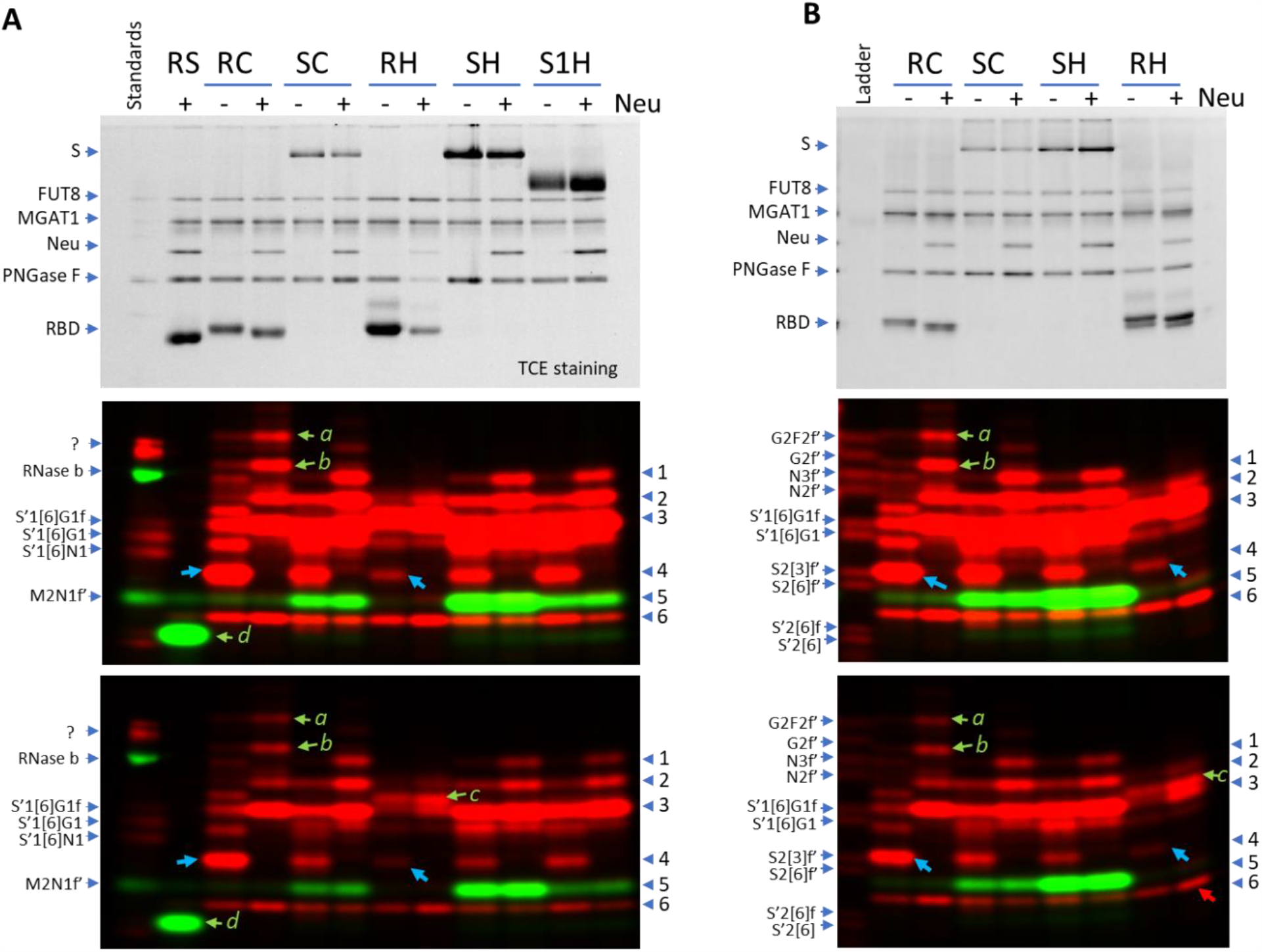
Fingerprinting N-glycans released from various SARS2 Spike proteins with ST6Gal1 (red) and FUT8 (green). N-glycans of various recombinant spike proteins released by PNGase F were labeled by ST6Gal1 and FUT8 with (+) or without (-) pretreatment of a neuraminidase (Neu). Labeled samples were separated on 17% SDS gel and imaged with regular protein imaging (upper panels) or fluorescent imaging (middle and lower panel in different contrasts). Bands in red are complex or hybrid glycans labeled by ST6Gal1 (via the incorporation of Cy5-conjugated sialic acid). Bands in green are oligo-mannose glycans labeled by FUT8 (via the incorporation of AlexaFluor 555-conjugated Fucose). For FUT8 labeling, MGAT1/UDP-GlcNAc were also included in the labeling mix to convert the oligo-mannose glycans to the substrates for FUT8. Labeled glycans from Ribonuclease B, S’1[6]N1, S’1[6]G1 and S’1[6]G1f were run as references in (A) and a glycan ladder with 10 labeled standard glycans was run in (B). RS, <u>R</u>BD domain of SARS2 Spike protein expressed in <u>S</u>f21 cells; RC, <u>R</u>BD domain of SARS2 Spike protein expressed in <u>C</u>HO cells; RH, <u>R</u>BD domain of SARS2 Spike protein expressed in HEK293 cells; SC, whole SARS2 Spike protein expressed in <u>C</u>HO cells; S1H, SARS2 <u>S1</u> protein expressed in <u>H</u>EK293 cells; SH, SARS2 <u>S</u>pike protein expressed in <u>H</u>EK293 cells.

Several common bands were observed (band 1 to 6) in Fig. 3. Band 1 and 2 were mainly found in neuraminidase treated SC, SH and S1H at the position around N3f’ and N2f’ (Fig. 3B). Since labeling through ST6Gal1 also contributes a sialic acid and therefore makes a labeled glycan move much faster, band 1 and band 2 could be due to labeling on highly branched complexed glycans, such as tetra- and tri-antennary complex glycans. Band 1 was mainly observed in desialylated SC, SH and S1H, suggesting that the glycan was initially sialylated. Band 2 was observed in SC, SH and S1H samples before and after desialylation, but with great signal increase upon desialylation, suggesting that the glycan was initially largely sialylated. Band 3 was prominent in all SH and S1H samples and in desialylated samples of RC and SC and had the same mobility of S’1[6]G1f (Fig. 3B). Since band 3 and the reference glycan S’1[6]G1f had the same labeling (both labeled on α,6-linked sialic acid with Cy5) and same mobility, band 3 likely had the same structure as S’1[6]G1f and was the labeling product of G2f (Fig. 2E). The fact that band 3 was much weaker in RC and SC than in desialylated RC and SC samples suggests that the glycan was initially sialylated in these samples. Opposite to band 3, band 4 around the position of S2[3]f’ had strong presence in RC and SC but not in the desialylated RC and SC samples, suggesting that band 4 was due to the labeling of a partially sialylated glycan that was converted to band 3 when desialyation occurred before labeling. Band 5 was likely due to the labeling of oligomannose M3 (known as Man5) as the band had same mobility of the reference glycan M2N1f’ (Fig. 3A). Band 6 had almost equal intensity in all samples and did not respond to *C*.*p* Neuraminidase treatment. The fast mobility of band 6 suggests that it is highly sialylated, but its unresponsiveness to neuraminidase treatment suggests the opposite. The nature of band 6 remains to be investigated.

Most of the common bands displayed great variation among the samples. For examples, band 4 was the most abundant in RC but almost at negligible level in RH (blue arrows in Fig 3); band 5 was the most abundant in SH but almost completely lacking in RH. Surprisingly, some bands were only found in one sample but not the others, such as band *a, b, c* and *d* (Fig. 3A). Band *a* and *b* in RC had slow mobility and responded to neuraminidase treatment, suggesting that they had highly complexed structures and were initially sialylated. Band *c* in RH was just above the position of S’1[6]G1f, suggesting that it might be S’1[6]G1fn that contains a bisecting GlcNAc (Fig, 2B and 2C). Band *d* labeled by FUT8 was found only in RS and had faster mobility than M2N1f’, suggesting that it be M1N1f’ (labeled product of Man3), in consistent to the notion that Man3 is a main glycan expressed in insect cells ^25^. Additional enzymatic conversion of band *d* with B4GalT1 and ST6Gal1 further confirmed the identity of band *d* (Supplemental Fig. 3).

### N-Glycan Fingerprinting Study of SARS-2 Spike proteins with ST3Gal6

Both ST6Gal1 and ST3Gal6 are known to sialylate the Galβ1,4GlcNAc structure on glycoproteins ^24^. When a same set of the SARS-2 spike protein samples were probed with ST3Gal6, similar but distinctive glycan fingerprints were observed (Fig. 4). It seems that the entire band pattern revealed by ST3Gal6 was upshifted from that of ST6Gal1. For examples, the bands 1’, 2’, 3’, and 6’ in ST3Gal6 labeled SH sample corresponded well with the bands 1, 2, 3, and 6 in ST6Gal1 labeled SH sample; similar to band 6, band 6’ was found across all lanes; similar to the relative positioning of band *b* and band 3 revealed by ST6Gal1, band *b’* labeled by ST3Gal6 was slightly upshifted from band 3’. The upshift observed of the bands revealed by ST3Gal6 from those by ST6Gal1 is likely due to that glycans with α,3-linked sialic acid had slower mobility than corresponding glycans with α,6-linked sialic acid (Fig. 2).

**Figure 4.**
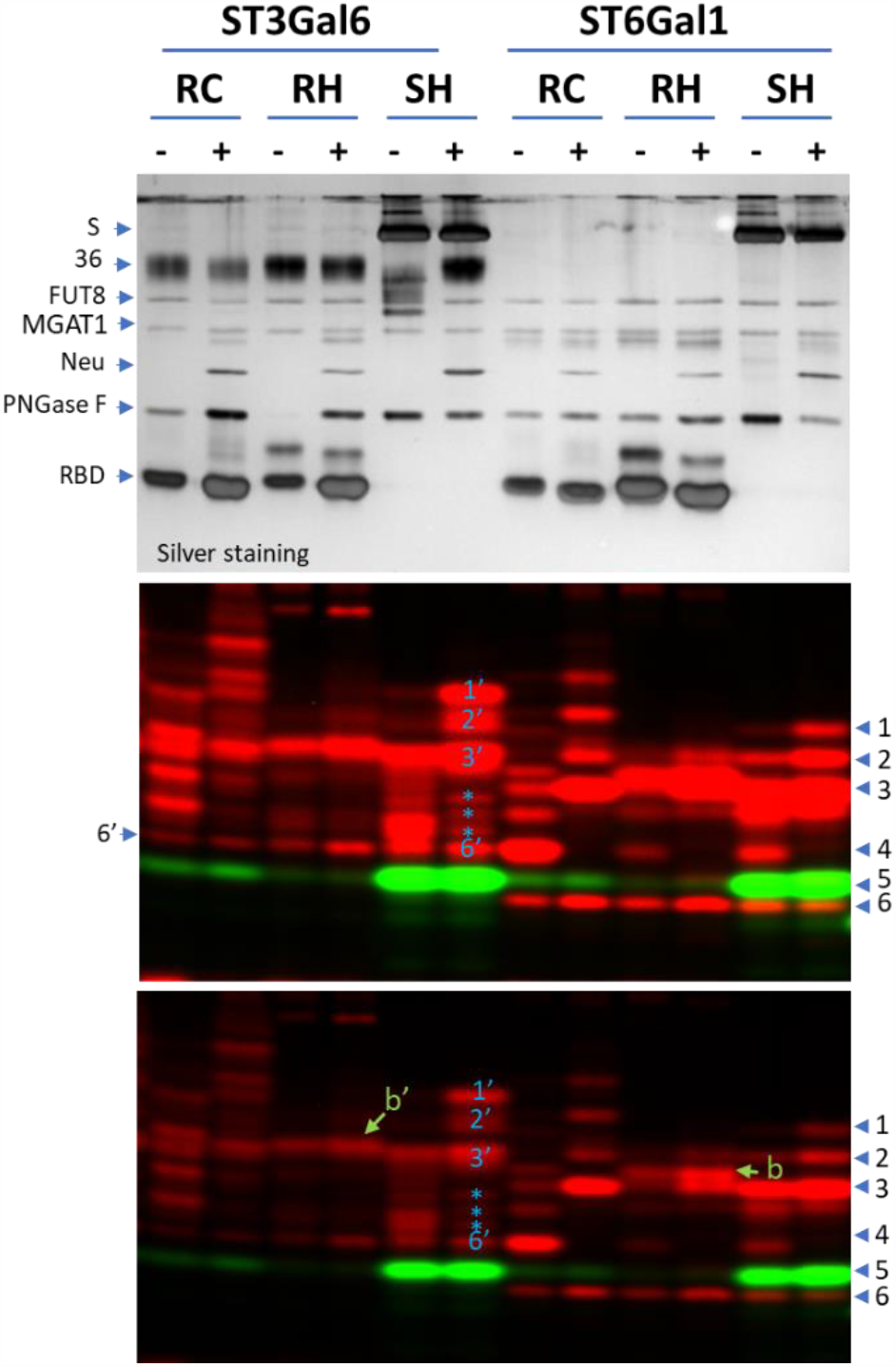
Fingerprinting by ST3Gal6 and ST6Gal1 on various SARS2 Spike proteins. Glycans of various recombinant SARS2 spike proteins released by PNGase F with (+) or without (-) neuraminidase pretreatment were labeled by ST3Gal6 or ST6Gal1 together with FUT8 and separated on 17% SDS gel. The gel was then imaged with silver staining (upper panel) or fluorescent imaging (middle and lower panel in different contrasts). Bands in red are complex or hybrid glycans labeled by ST6Gal1 or ST3Gal6 (via the incorporation of Cy5-conjugated sialic acid). Bands in green are oligo-mannose glycans labeled by FUT8 (via the incorporation of AlexaFluor 555-conjugated Fucose). For oligo-mannose labeling, MGAT1 and UDP-GlcNAc were also included in the labeling mix to convert the glycans to the substrates for FUT8. RC, <u>R</u>BD domain of the SARS2 Spike protein expressed in <u>C</u>HO cells; RH, <u>R</u>BD domain of the SARS2 Spike protein expressed in <u>H</u>EK293 cells; SH, whole SARS2 <u>S</u>pike protein expressed in <u>H</u>EK293 cells.

While there was similarity between the fingerprintings revealed by the two enzymes, ST3Gal6 labeling also revealed some unique bands. For example, bands marked with asterisks in SH revealed by ST3Gal6 had no corresponding bands in SH revealed by ST6Gal1. This difference on band patterning suggests that ST3Gal6 and ST6Gal1 have overlapping but distinctive substrate preferences.

### Finger printing of O-Glycans of SARS-2 Spike proteins with ST3Gal1

O-glycosylation of the RBD domain of SARS-Cov-2 spike protein has been reported previously ^18, 26^. Here, we investigated whether finger printing technique could be applied to study O-glycans as well. One challenging issue for finger printing O-glycans is that there is no enzyme (corresponding to PNGase F for removal of N-glycans) that allows for wide removal of O-glycans. Currently, *E. faecalis* O-glycoidase (Endo EF) is used to remove Core-1 type O-glycan ^27^. Another challenging issue is that O-glycans are usually smaller than N-glycans and rather difficult to be separated from the donor substrates of various glycosyltransferases in SDS-gel.

To overcome the above challenges, we first directly probed O-glycans on the RBD proteins using O-glycan specific ST3Gal1 as previously described ^20^ (Fig. 5A). Indeed, O-glycans were detected on all RBD proteins investigated, but with different levels of sialylation and sensitivity to Endo EF treatment (Fig. 5B). The labeling on the RBD proteins expressed in CHO cells, Sf21 cells and Tn cells were removed by Endo EF treatment, suggesting that these samples contain Core-1 O-glycan. To understand the difference among the O-glycans on RBD proteins expressed in HEK293 cells, the RBD protein expressed in Sf21 cells was first pretreated with GCNT1 ^28^, a GlcNAc transferase that converts Core-1 O-glycan to Core-2 O-glycan, or together with B4GalT1 ^29^ that can extend the GlcNAc residue on Core-2 O-glycan, and then labeled with Cy5-Sialic acid by ST3Gal1, digested with trypsin, and finally separated in SDS-PAGE (Fig. 5C). Indeed, trypsin digestion resulted in two labeled glycopeptides in all samples, but with different mobility. The mobility of the two peptides were shifted by GCNT1 and B4GalT1 and finally matched to that of the peptides of RBD protein expressed in HEK293 cells. These results suggest that Oglycans on the RBD protein expressed in HEK293 are extended Core-2 type O-glycan, whereas the O-glycan on the RBD protein expressed in Sf21 cells is Core-1 type.

**Fig. 5.**
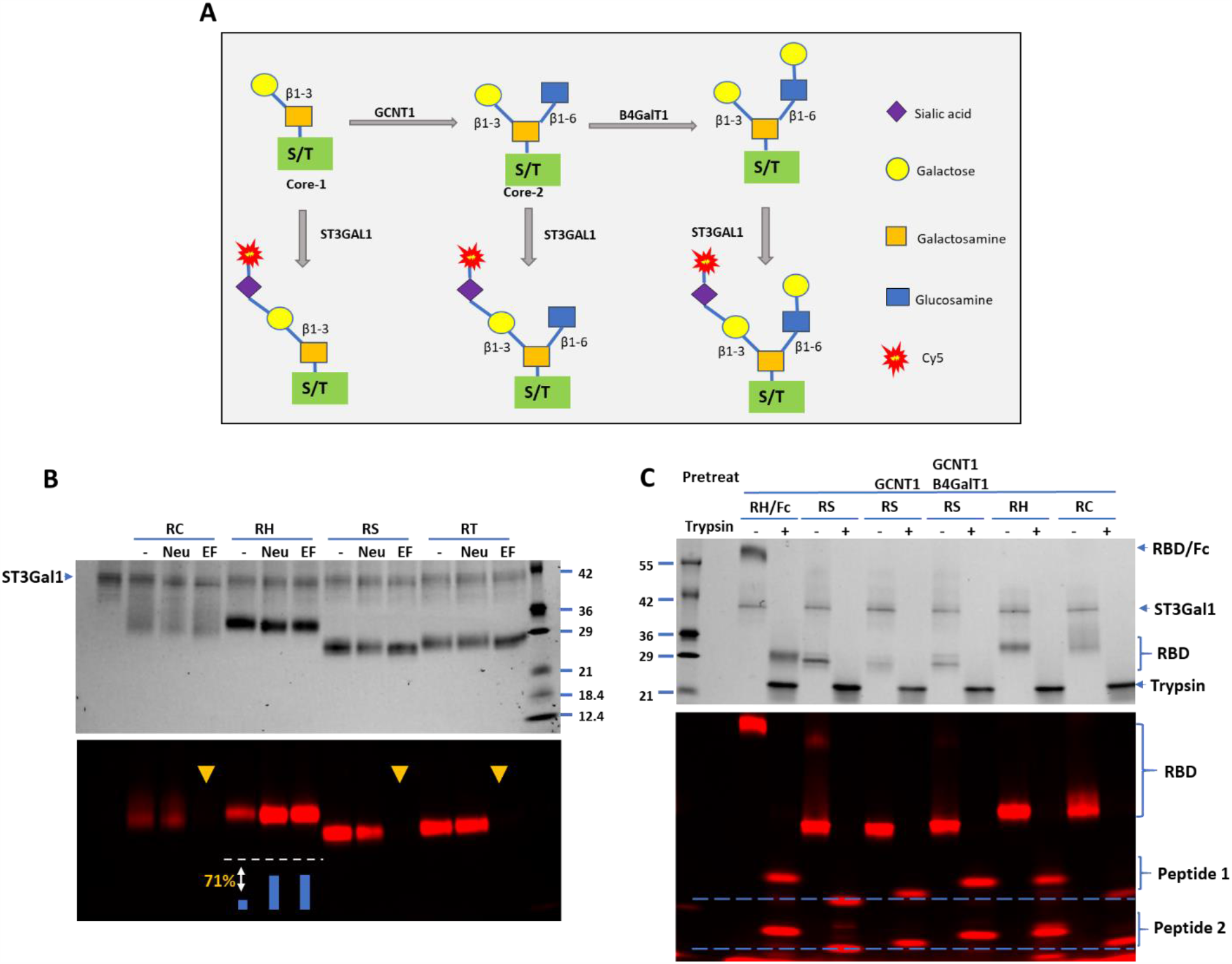
O-Glycan of the RBD domain of SARS-CoV-2 Spike protein varies depending on the host cells. **A**) Strategy for O-glycan conversion and labeling. **B)** RBD expressed in CHO (RC), HEK293 (RH), Sf21(RS) and Tn (RT) cells were probed for O-glycans with CMP-Cy5-Sialic acid and ST3Gal1. The proteins were directly labeled or labeled after pretreatment with *C. perfringens* neuraminidase alone (Neu) or in combination with Endo EF (EF). Densitometry analysis indicated that neuraminidase treatment resulted in 71% increase on labeling of the RBD protein expressed in HEK293 cells. **C**) RBD protein expressed in Sf21 cells (RS) pretreated with GCNT1 alone or together with B4GalT1 was labeled with ST3Gal1 and then digested with trypsin. RBD protein fused to IgG Fc domain expressed in HEK293 cells (RH/Fc), RH and RC samples were also labeled and digested by trypsin and run along RS samples.

## Discussion

In this report, we have established a novel method for N-glycan fingerprinting based on enzymatic fluorescent glycan labeling and electrophoresis. As a method of fingerprinting, the overall glycan patterning rather than individual glycan species is the focus. This method can serve as a quick and inexpensive way to interrogate if different batches of glycoproteins are consistent on glycosylation and screen out samples of abnormal glycosylation. Herein, this strategy is demonstrated on several SARS2 spike proteins, in which N-glycans are released and enzymatically labeled with fluorophore-conjugated sialic acid and fucose and finally separated on SDS-PAGE. Although the strategy does not allow site specific and detailed structural glycan analysis, it does offer some major advantages. First, it is simple, convenient, and much more affordable. Second, the data acquired are visually informative and therefore rather easy to interpret. Third, multiple samples can be processed simultaneously, therefore it is highly efficient. Fourth, the signal intensity is directly related to the abundance of a glycan species and therefore is quantitative ^21^. While this method only reveals the substrate glycans of the labeling enzyme, and glycans that are not recognized by the labeling enzyme remain to be undetected, which could be advantageous as well when only specific glycans are being examined.

Recognizing that fingerprinting is focused on general band patterning, it is always beneficial if the identities of individual bands on a gel can be determined. To this purpose, glycan standards and glycan ladders can be run along with samples to serve as references. By comparing the mobility of labeled bands to the mobility of the reference glycans, the identity of a labeled band maybe determined. Besides, it is found that the addition of a linkage specific monosaccharide changes the mobility of a glycan at relatively constant rate (Fig. 2). The knowledge of mobility shift caused by the addition of certain monosaccharides also allows us to deduce the identities of labeled bands. More specifically, the addition of a neutral monosaccharide slows down a glycan and the addition of a negatively charged sialic acid increases the mobility of a glycan. Particularly, a bisecting GlcNAc causes about half of the mobility shift caused by a β,6-linked GlcNAc and an α,6-linked sialic acid causes slightly more mobility shift than an α,3-linked sialic acid.

Our data suggest that the RBD of the SARS2 spike protein expressed in HEK293 cells mainly contains complex glycans, which is consistent with the reports of Watanabe, *et al* ^6^. Our data also suggests that bisecting GlcNAc may exist on the RBD portion, and oligomannose glycans may mainly exist on S protein except the RBD portion when expressed in HEK293 cells. Our data also indicate that the glycans of S proteins expressed in insect cells and HEK293 cells are completely different. Besides, using similar technique, we found that insect cells expressed RBD protein contains Core-1 O-glycan and HEK293 cells expressed RBD protein may contain extended Core-2 O-Glycan. Altogether, our study provides evidence to support that host cell determines the types of glycans attached to the spike proteins of SARS2, which further implies that the glycosylation pattern of the spike protein of SARS2 from COVID-19 patients could be different as well.

## Material and methods

Different recombinant SARS-CoV-2 Spike RBD proteins expressed in HEK293 cells, Tn5 insect cells, CHO cells, full length recombinant SARS-CoV-2 Spike proteins expressed in HEK293 cells and CHO cells, and recombinant SARS-CoV-2 Spike S1 subunit protein expressed in HEK293 cells were from Bio-Techne. Recombinant human ST6Gal1, FUT8, B4GalT1, MGAT1, ST3Gal6, FUT9, GCNT1, ST3Gal4 and ST3Gal3, and *C. perfringens* neuraminidase and *F. meningosepticum* PNGase, *E. faecalis* O-Glycosidase, CMP-Cy5-Siallic acid, GDP-AlexaFluor555-Fucose were from Bio-Techne. IgG glycan G0, G1F and G0F were from Dextra Labs. Trypsin was from Sigma Aldrich.

### Releasing and Labeling of the N-Glycans of Spike Proteins

To release N-glycans, 5 μg of a spike protein was mixed with 0.2 μg PNGase F and supplemented with labeling buffer (25 mM Tris pH 7.5, 10 mM MnCl_2_) to 20 μl and then incubated at 37° C for 30 minutes. For desialylation, an additional 0.2 μg *C*.*p*. neuraminidase was also added into the reaction mixture. The above mixture was then heated at 95°C for two minutes to inactivate the enzymes. Labeling mixture contained 0.5 μg of a sialyltransferase together with 0.4 nmol of CMP-Cy5-Sialic Acid supplemented with labeling buffer to 10 μl. In the case for labeling oligomannose, additional 0.5 μg of FUT8 together with 0.4 nmol of GDP-AlexaFluor555-Fuc and 0.5 μg of MGAT1 together with 10 nmol of UDP-GlcNAc were also added into the labeling mixture. The labeling mixture was then added into the reaction mixture and incubate at 37° C for 1 to 2 hours or overnight at room temperature.

### Labeling Glycan Standards and Building Glycan Ladder

For labeling a glycan with Cy5-Sialic Acid, 1 μg of the standard was mixed with 1 μg of ST6Gal1 and 1 nmol of CMP-Cy5-Sialic Acid together with 0.5 μg B4GalT1 and 10 nmol of UDP-Gal supplemented with labeling buffer to 20 μl and the mixture was incubated at 37° C for 2 hours or at room temperature for overnight. For labeling a glycan standard with Cy5-Fucose, 2 μg of the standard was mixed with 1 μg of FUT8 and 2 nmol of GDP-Cy5-Fucose supplemented with labeling buffer to 20 μl and the mixture was incubated at 37° C for 2 hours or at room temperature for overnight. For building a glycan ladder based on Cy5-Fucose labeled glycan standard, 200 ng of the above labeled glycan was extended with additional one or more of 0.5 μg each of the glycosyltransfeases including MGAT3, MGAT5, B4GalT1, FUT9, ST3Gal6 and ST6Gal1 together with their donor substrates at 37° C for 2 hours or overnight at room temperature or whenever the reactions were completed. The reactions were then stopped by heating at 95° C for 2 minutes. Glycan ladder was built by mixing equal amounts of the above extended labeled glycans.

### Glycan Electrophoresis and Imaging

All labeled samples including glycan standards were separated on 15% or 17% sodium dodecyl sulfate–polyacrylamide gel electrophoresis (SDS-PAGE) at 20 volts/cm. After separation, all gels were imaged using a FluorChem M imager (ProteinSimple, Bio-techne). For imaging protein contents, the gel was also imaged with traditional methods such as silver staining or trichloroethanol (TCE) staining.

## Acknowledgements

We would like to thank all colleagues who have made contribution to this work through product development

