## Supplemental Fig. 1, 2 and 3 for "Fluorescent Glycan Fingerprinting of SARS2 Spike Proteins"

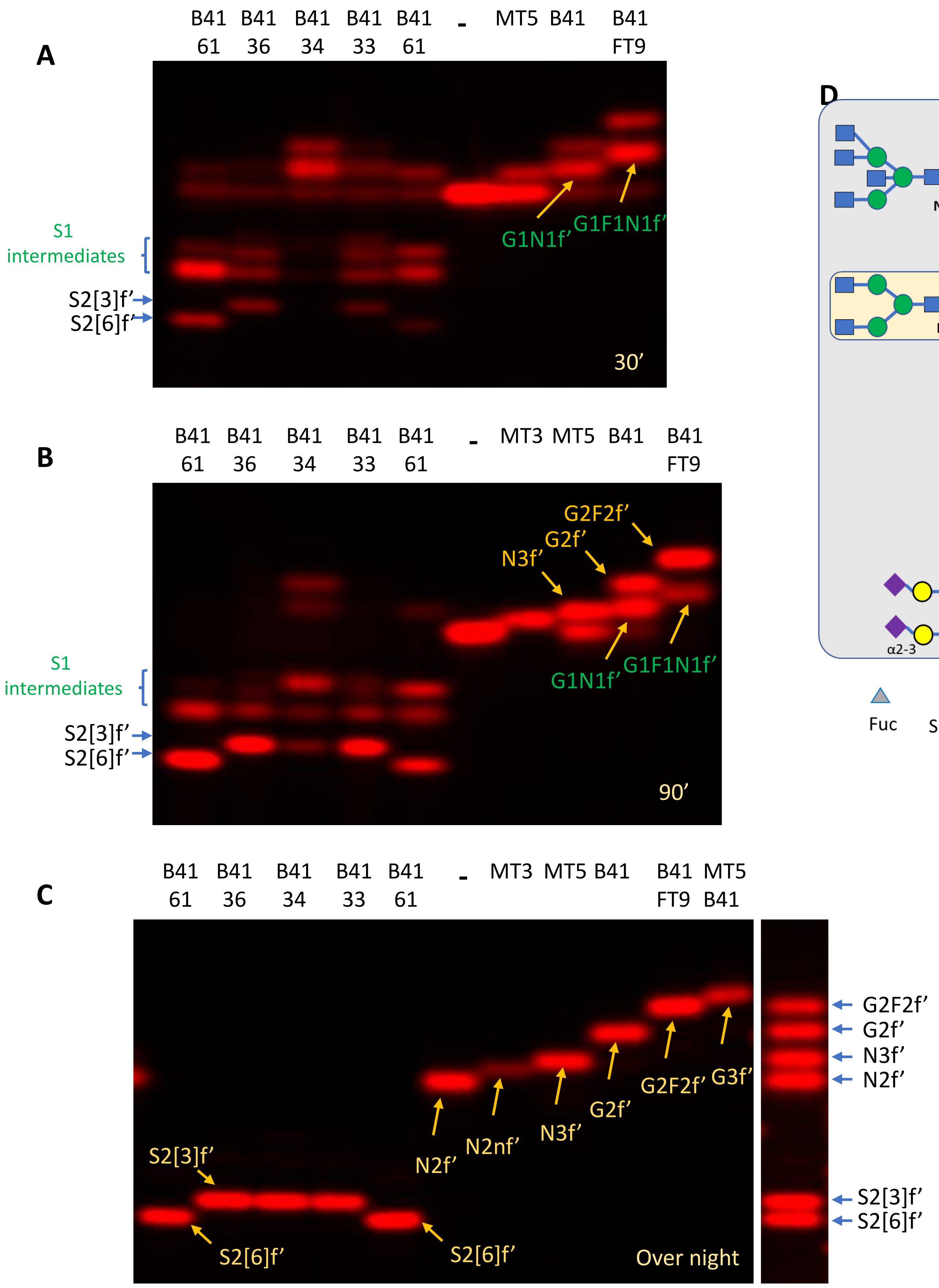


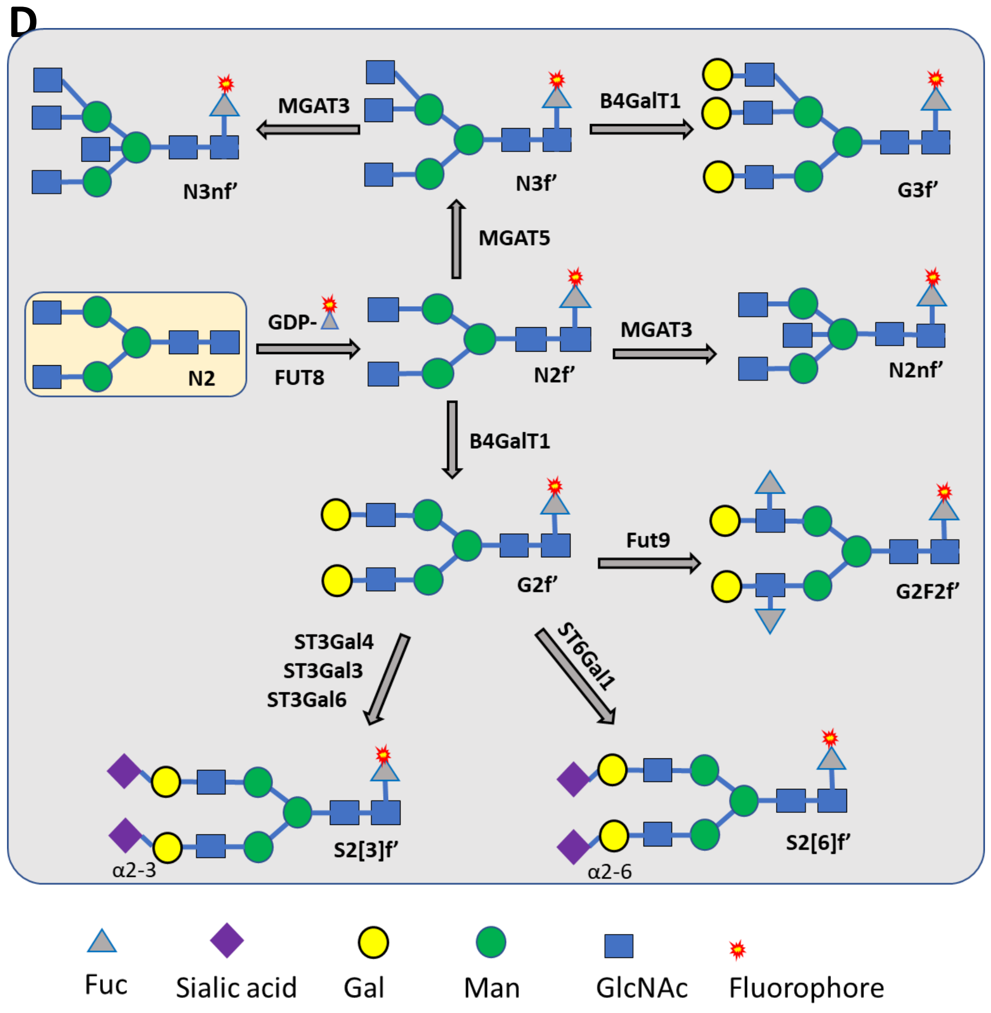


**Supplemental Fig. 1 Intermediates During the Generation of the Final Labeled Glycans .** All glycans were enzymatically extended from N2f' that was labeled at the core-fucose by Fut8 with GDP-Cy5-Fuc. Enzymatic reactions were incubated at room temperature for 30 minutes (**A**), 90 minutes (**B**) and overnight (**C**) and separating on 17% gel. Synthesis of N2nf' and G3f' were not started initially and were only introduced in (**B**) and (**C**) respectively. Enzymes used for the conversion are indicated above the images. B41, B4GalT1; 61, ST6Gal1; 36, ST3Gal6; 34, ST3Gal4; 61, ST6Gal1; MT3, MGAT3; MT5, MGAT5, F9, Fut9. Short names of glycans and schemes for enzymatic generation of these glycans are indicated in (**D**). For rules of naming these glycans please refer to Fig.1.

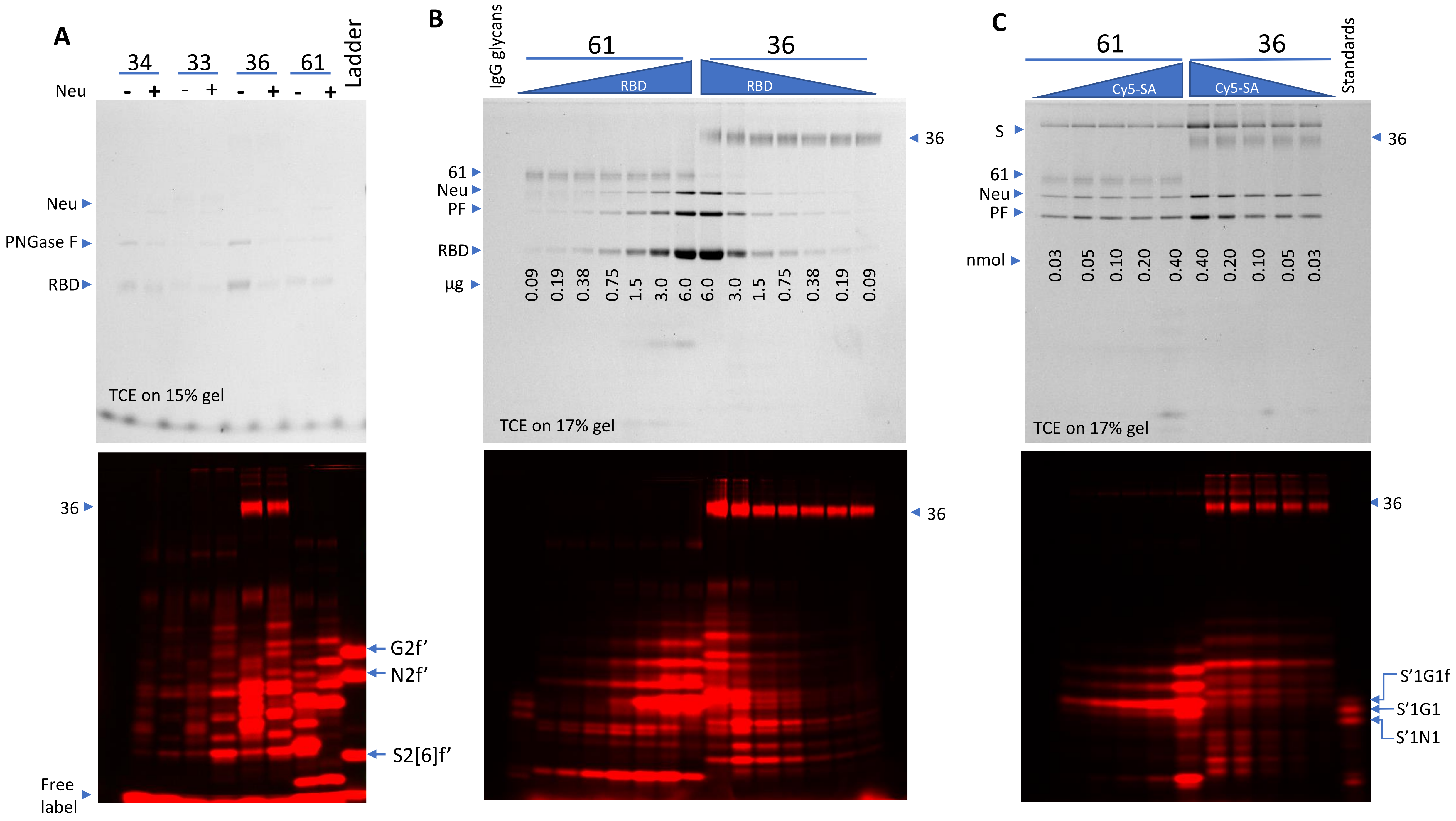


**Supplemental Fig. 2 Selection of ST6Gal1 and ST3Gal6 for glycan finger printing study and their substrate concentration optimization.** In all cases, labeling reactions were proceeded for 2 hours at 37°C and then separated on indicted SDS-gel, and, imaged by TCE staining (top panels) and fluorescent imaging (low panels). 33, ST3Gal3; 34, ST3Gal4; 36, ST3Gal6; 61, ST6Gal1. ST3Gal6 showed self-labeling as indicated. **(A)** Glycans from SARS2 RBD protein expressed in CHO cells were released by PNGase F and labeled by various sialyltransferases with CMP-Cy5-Sialic acid. Samples were also labeled with (+) or without (-) neuraminidase (Neu) pretreatment. The glycan ladder contained G2f', N2f' and S2[6]f'. **B**) Variable amounts of the RBD protein were labeled by ST6Gal1 or ST3Gal6. **C**) Variable amounts of CMP-Cy5-Sialic acid were used to label the S protein expressed in CHO cells by ST6Gal1 or ST3Gal6.

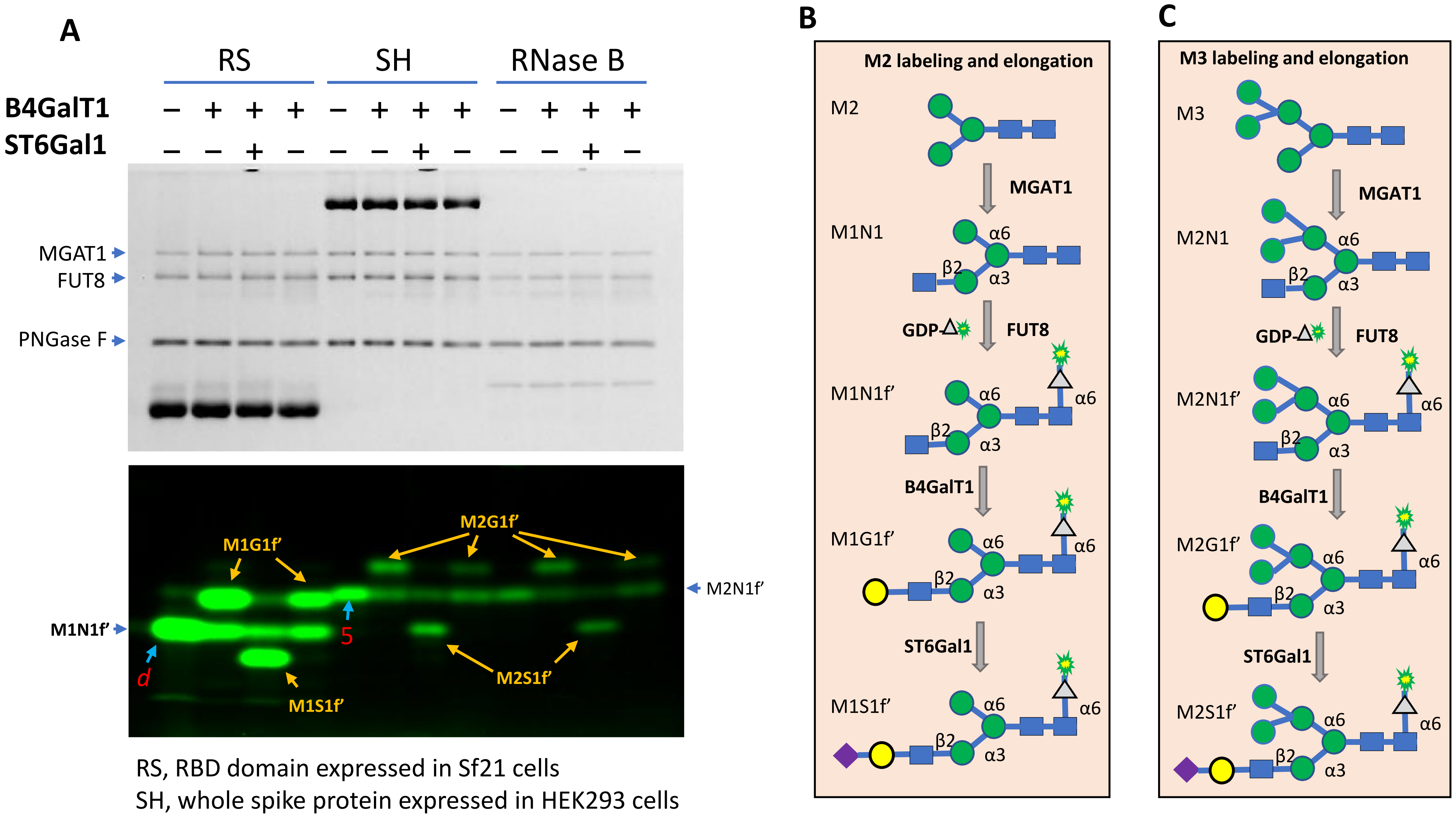


RS, RBD domain expressed in Sf21 cells
SH, whole spike protein expressed in HEK293 cells

**Supplemental Fig 3. Mobility shift of band *d* and band 5 of Fig. 3 by enzymatic modification**. Glycans of the recombinant RBD expressed in Sf21 cells (RS) and whole spike protein expressed in HEK293 cells (SH) along with that of RNase B released by PNGase F were first labeled by FUT8 with GDP-Cy3-fucose together with MGAT1 and then further treated with B4GalT1 and ST6Gal1 (indicated with + and – signs). RNase B known to contain Man5 (M3) was used as a control. Samples were run on 17% gel visualized by TCE imaging (upper panel) and fluorescent imaging (lower panel) (**A**). The scheme for labeling and molecular conversion of MAN3 (M2) is shown in (**B**). The scheme for labeling and molecular conversion of M3 is shown in (**C**). The labeled bands from SH and RNase B are at same the position, suggesting that the glycan released from SH is indeed M3. The spacing between the labeled band of RS and SH suggests that the glycan released from RS is M2. The shifts on the labeled bands from RS and SH caused by B4Galt 1 and ST6Gal1 are the same, further confirming that the glycan released from RS is M2.
